# Copy number variations with adaptive potential in caribou (*Rangifer tarandus*): genome architecture and new annotated genome assembly

**DOI:** 10.1101/2021.07.22.453386

**Authors:** Julien Prunier, Alexandra Carrier, Isabelle Gilbert, William Poisson, Vicky Albert, Joëlle Taillon, Vincent Bourret, Steeve D. Côté, Arnaud Droit, Claude Robert

**Affiliations:** Genomic center - Laval University; Laval University; Ministere des Forets, Faune et Parcs du Quebec; Ministere Foret, Faune et Parcs du Quebec

**Keywords:** Genome assembly, transcriptome assembly, copy number variation, *Rangifer tarandus*, genome annotation

## Abstract

**Background:** *Rangifer tarandus* (caribou and reindeer) has experienced recent drastic population size reductions throughout its circumpolar distribution. In efforts aimed at preserving caribou in North America and reindeer in Eurasia, genetic diversity conservation is of utmost importance, particularly the adaptive genetic diversity. To facilitate genomic studies of the caribou population, we improved genome assembly and annotation by combining long-read, linked-read and RNA sequencing technologies. As copy number variations (CNVs) are known to impact phenotype and are therefore likely to play a key role in adaptation, we investigated CNVs among the genomes of individuals representing three ecotypes of caribou (migratory, boreal and mountain).

**Results:** Using *de novo* transcriptome assembly and similarity with annotated human gene sequences, we identified 17,394 robust gene models embedded in a new highly contiguous genome assembly made of 13,994 scaffolds and presenting the highest N50 reported to date. A BUSCO analysis supported the high accuracy of this assembly, 90% of which being represented by only 131 scaffolds. Genome level comparisons with domestic ruminant species showed high synteny within this clade. A total of 1,698 large CNVs (length > 1kb) were identified, including 332 overlapping coding sequences annotated for functions related to immunity, musculoskeletal development or metabolism regulation and others. While the CNV distribution over the genome revealed 31 CNV hotspots, 43 large CNVs were particularly distinctive of the migratory and sedentary ecotypes and included genes annotated for functions related to cardiac development, fatty acid regulation, cold responses, locomotory behavior or environmental perception (hearing and sight), that can be related to the expected adaptations.

**Conclusions:** This work includes the first publicly available annotation of the *Rangifer tarandu*s genome and the first genome assembly allowing genome architecture analyses. This robust annotation based on truly expressed sequences showed a distribution overlapping many CNVs that are promising candidates given the annotations supporting their involvement in adaptation. This new highly contiguous assembly will allow relative localization of genetic variations and features and will be a valuable resource for molecular tool development and genomic studies aimed at describing and preserving this species.

## Background

The genome architecture of adaptation is an important factor contributing to the evolution of a species (Feder and Nosil 2010; Yeaman 2013). Among the genetic variations potentially related to adaptation, structural variations (SVs), including copy number variations (CNVs), have been associated with phenotypic variations and local adaptations (Wellenreuther et al. 2019; Mérot et al. 2020). Since the first large-scale screenings showing that CNVs in human genomes involve more nucleotides than single-nucleotide polymorphisms (Carson et al. 2006; Redon et al. 2006; Sebat et al. 2004; Itsara et al. 2010; Conrad et al. 2010), an increasing number of studies even suggested that CNVs have a greater impact than SNPs on phenotypic variations (de Smith et al. 2008) and consequently on adaptation (Mérot et al. 2020).

CNVs are usually defined as DNA segments longer than 1 kb occurring in various copy numbers within a species, such copies presenting an identity exceeding 90% (Redon et al. 2006; Sebat et al. 2004; Feuk, Carson, and Scherer 2006; Freeman 2006). CNVs do not arise from transposable elements (Freeman 2006) but from a variety of mechanisms including non-allelic homologous recombination (NAHR), non-homologous end joining (NHEJ), single-strand annealing, breakage-fusion-bridge cycle, or replicative non-homologous DNA repair (Hastings et al. 2009; Gu, Zhang, and Lupski 2008; Lovett 2004). A majority of these mechanisms are related to the occurrence of low-copy repeats (LCRs or tandem repeats) which occur throughout the genome and present nucleotide sequence identity exceeding 95%. As a result, CNVs tend to cluster into hotspots found in the surroundings of these LCRs (Hastings et al. 2009).

Structural variations may appear *de novo* in somatic tissues where they can cause pathologies such as cancers for instance, or in the germline in which case they may be transmitted to the next generation and result in heritable phenotypic variations (Gu, Zhang, and Lupski 2008). The mutation rate for CNVs has been estimated at ∼1×10-4, which is higher than the SNP mutation rate (Lupski 2007). They may impact phenotype through the gene dosage effect, i.e. a gene copy number variation resulting in gene expression variation that affects the phenotype (Gamazon and Stranger 2015; Perry et al. 2007), but they can also trigger sequence disruption (gene sequence truncation) or fusion, or even have position effects (Lupski and Stankiewicz 2005). As a result, purifying selection may select against CNV encompassing genes, particularly deletions that are less likely tolerated than gene duplication (Brewer et al. 1999; Conrad 2006).

Copy Number Variations present several interesting characteristics regarding the genomic architecture of adaptation that can contribute to species evolution. Large CNV sequences may span more than one gene, and such gene clusters may collectively have an impact on phenotype, for example, nematode resistance in maize (Cook et al. 2012). In addition, since CNVs tend to cluster into genomic hotspots (Hastings et al. 2009), they may be inherited as clusters of locally adaptive loci and thus confer an adaptive advantage (Yeaman 2013). Finally, CNVs may prevent recombination and thus promote large genomic islands of divergence favoring the apparition and persistence of adaptations to local conditions (Puig Giribets et al. 2019; Tigano et al., n.d.).

CNVs have been investigated in a number of domestic mammal species including cattle (Fadista et al. 2010; Hu et al. 2020), swine (Wang et al. 2013), horses (Wang et al. 2014), sheep (Fontanesi 2011) and goats (Fontanesi 2010). These early genome-wide CNV studies revealed relatively few CNVs (37 to 368) per genome with length averaging 127 kbp to 10.7 Mbp due to the low resolution inherent in detection methods based on array comparative genomic hybridization (aCGH) or SNP chips (Clop, Vidal, and Amills 2012). Nevertheless, comparison of bovine, caprine and ovine large CNV maps shows substantial overlap (Clop, Vidal, and Amills 2012; Fontanesi et al. 2011), which is attributed to conservation of segmental duplications in these regions, promoting recurrent CNVs through NAHR rather than CNVs inherited by descent (Clop, Vidal, and Amills 2012). As observed in the human genome, genes included in livestock CNVs tend to be annotated for functions related to immunity, sensory perception, amongst others (Clop, Vidal, and Amills 2012). More recent results obtained from higher resolution techniques and more exhaustive genome scans have corroborated such results in horses (Schurink et al. 2018) and goats (Dong et al. 2015; Genova et al. 2018) and revealed pigs CNVs that span genes annotated for functions related to metabolism and olfactory perception (Paudel et al. 2015). In addition, CNVs are involved in the between-race phenotypic diversity in dogs, including height for instance (Serres-Armero et al. 2021).

However, CNVs remain scarcely investigated at the genome scale in comparison with SNPs, particularly in wild species. This is largely due to the challenges inherent in CNV discovery at the genome level which has long relied on aCGH, now replaced by read-depth (coverage) and read-distribution-based approaches made possible by the advent of second-generation sequencing (Alkan, Coe, and Eichler 2011). In both cases, high-quality genome assembly is required, which is often lacking for undomesticated species, although there have been exceptions (Prunier, Caron, and MacKay 2017, for a gene-based aCGH approach).

In the present study, we investigated CNVs in caribou (*Rangifer tarandus*), a wild ruminant in North America. Several populations of this emblematic mammalian species with a circumpolar distribution have declined in the last decades and are endangered by climate change and human activities (Festa-Bianchet et al. 2011; Vors and Boyce 2009). Caribou in Northeastern America are divided into three major ecotypes: the migrating caribou, which spend the winter in the forest but calve and spend the summer in the tundra, the sedentary boreal caribou, which remain in the boreal forest all year and do not migrate, and the mountain caribou, which inhabit relatively low mountain tops (Mallory and Hillis 1998). This diversity of habitats exposes the species to a variety of selective pressures in terms of predation and parasites, competition with other ungulates, as well as varying forage composition (Mallory & Hillis 1998). For example, sedentary boreal caribou usually travel only a few kilometers while migrating caribou travel hundreds to thousands of kilometers annually (Mallory & Hillis 1998). In addition, migrating caribou are more prone to harassment from *Oestridae* parasitic flies that are increasingly active with increasing solar radiation in the toundra (Hagemoen and Reimers 2002) while sedentary boreal caribou are relatively spared in the shade of the boreal forest.

We report here a new *Rangifer tarandus* genome assembly based on long reads and linked reads to improve completeness and quality (Warren et al. 2017), which we annotated using RNAseq *de novo* assembly and gene annotation from other mammals. We detected CNVs using short-read sequencing from individuals representing the three ecotypes, expecting to find ecotype-specific CNVs involving genes with annotations likely related to the different ecological conditions of the three ecotypes. Our results provide support for genomics tool development and fine-scale genomic studies of caribou.

## Results

### An improved genome assembly for a wild ruminant

We used the following three strategies to obtain a high-quality contiguous assembly of the genome of a female caribou: long reads with PacBio SMRT cells, Illumina 2×150 bp linked reads from a Chromium 10X library, and Illumina 2×150 bp paired end sequencing of 400 bp inserts. PacBio SMRT cells yielded 7,534,419 high-quality long reads averaging 10,108 bp and representing an uncorrected coverage of 47X (assuming a genome size of 3 Gbp). Chromium 10X library sequencing using Illumina HiseqX yielded 2,140,002,320 linked reads of 150 bp representing a coverage of 107X. Finally, 813,953,740 short reads of 150 bp were obtained with Illumina sequencing, representing a coverage of 40X.

Assembling the long reads using Falcon (Chin et al. 2016) yielded a 2.52 Gbp genome assembly composed of 6,351 contigs (N50 = 501,648 bp, Fig. 1). This assembly accuracy was supported by the BUSCO analysis that found almost all mammalian conserved orthologous genes (C: 90.3%, F: 7%). Assembling linked reads with Supernova yielded 21,785 scaffolds for a total of 2.56 Gbp (N50 = 2,383,988 bp). Almost all mammal conserved orthologous genes were again found (C: 91.7%, F: 4.2%). The Falcon assembly was then scaffolded using the Supernova assembly and the resulting assembly was re-scaffolded using a public caribou genome assembly obtained using the DoveTail approach (Taylor et al. 2019).

**Figure 1.**
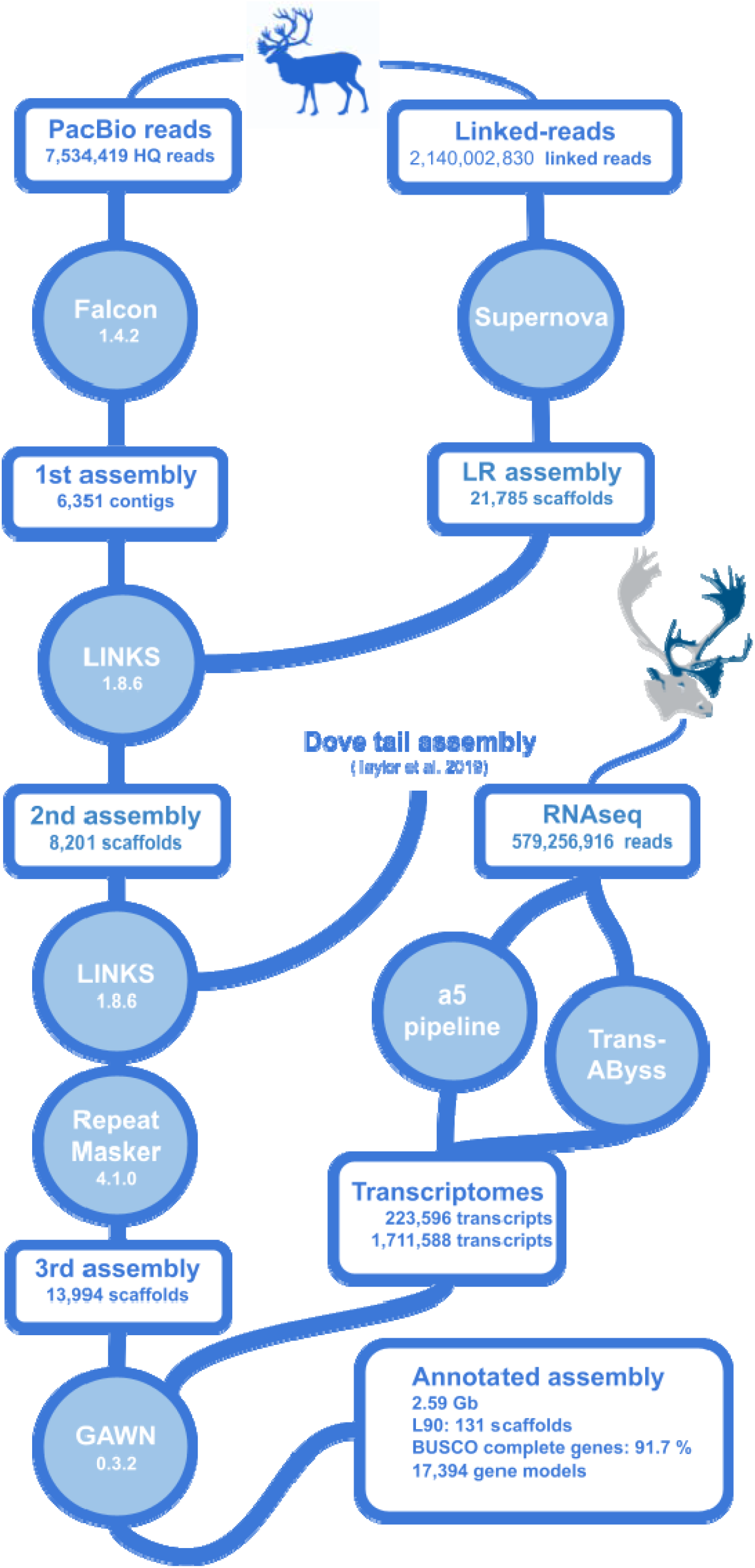
Caribou genome assembly and annotation pipeline

The final 2.59 Gbp assembly contained fewer and longer contigs and scaffolds than assemblies published so far for this species and thus represented a significant improvement (Fig. 2), particularly in terms of the number of scaffolds representing 90% of the assembly (L90) (Table 1). Using short reads assembled independently or to correct long reads did not improve the genome assembly in terms of contiguity (N50) or accuracy (BUSCO analysis).

**Table 1.** *Rangifer tarandus* genome assemblies published or obtained in this study.

| Publication | Total sequence length (Gbp) | Number of scaffolds | Scaffold N50 (Kbp) | L90 scaffold number | GC content | BUSCO analysis (total DB size)* |
| --- | --- | --- | --- | --- | --- | --- |
| Li et al. 2017 | 2.64 | 58,765 | 986 | -- | 41.2% | 92.6% (4104) |
| Taylor et al. 2019 | 2.21 | 4,699 | 11,765 | 289 | 41.4% | 93.1% (4104) |
| Weldenegodguad et al. 2020 | 2.66 | 23,450 | 5023 | -- | 41.4% | 92.9% (4104) |
| The present study | 2.59 | 13,994 | 29,299 | 131 | 41.5% | 91.7% (9226) |

\* Only the ratio of complete single-copy gene sequences are shown here; gene database size in parentheses

**Figure 2.**
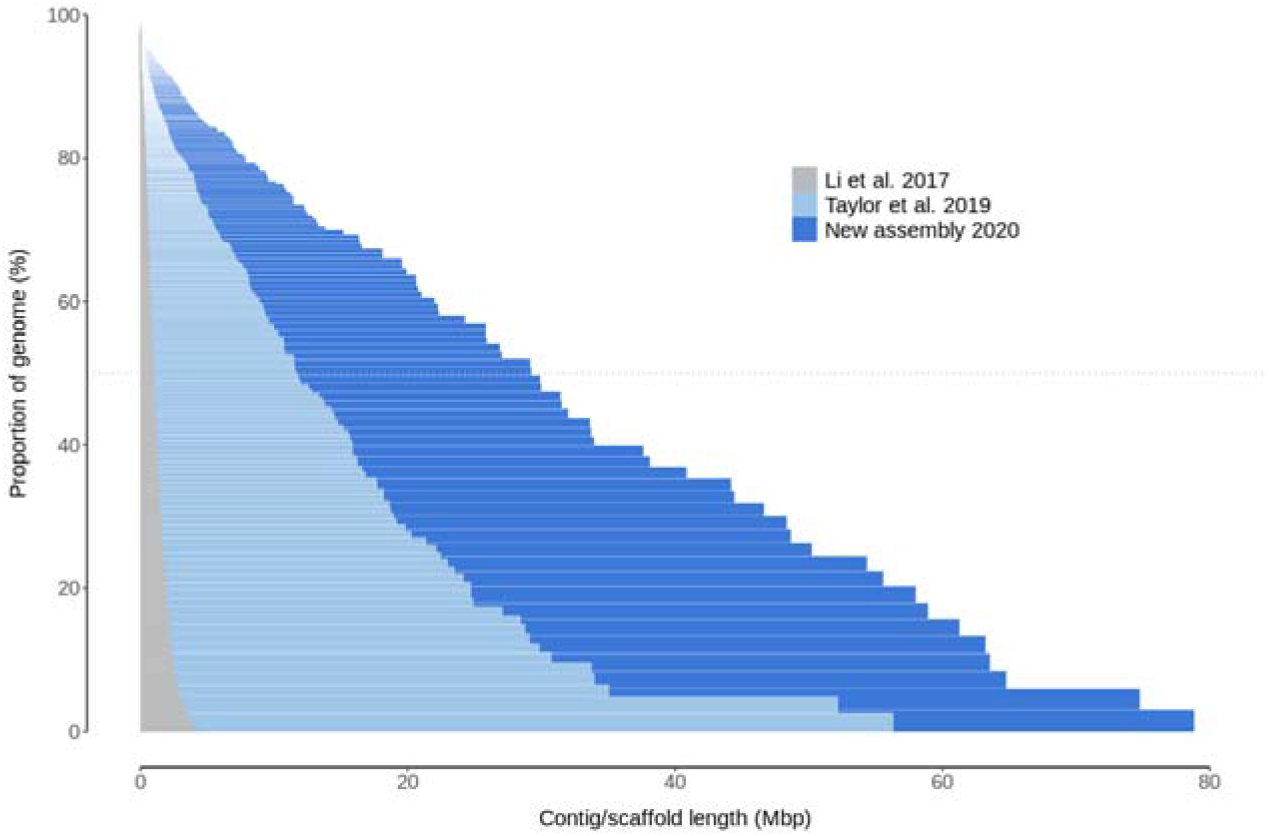
Scaffold length distributions in published *Rangifer tarandus* genome assemblies

### High synteny and phylogenetic clustering with other ruminant genomes

*Bos taurus* and *Capra hircus* genomes were compared on the basis of scaffold alignment with reference genomes using minimap2 (Li 2018) and visualization integrated into the JupiterPlot bioinformatic tool (Chu et al. https://github.com/JustinChu/JupiterPlot). Since representation was found to be the same for these ruminant genomes, only the comparison with *Bos taurus* is plotted in Figure 3. In both cases, a very high synteny was observed, although 18 crossing lines and bands indicated variations in DNA segment order and contiguity.

**Figure 3.**
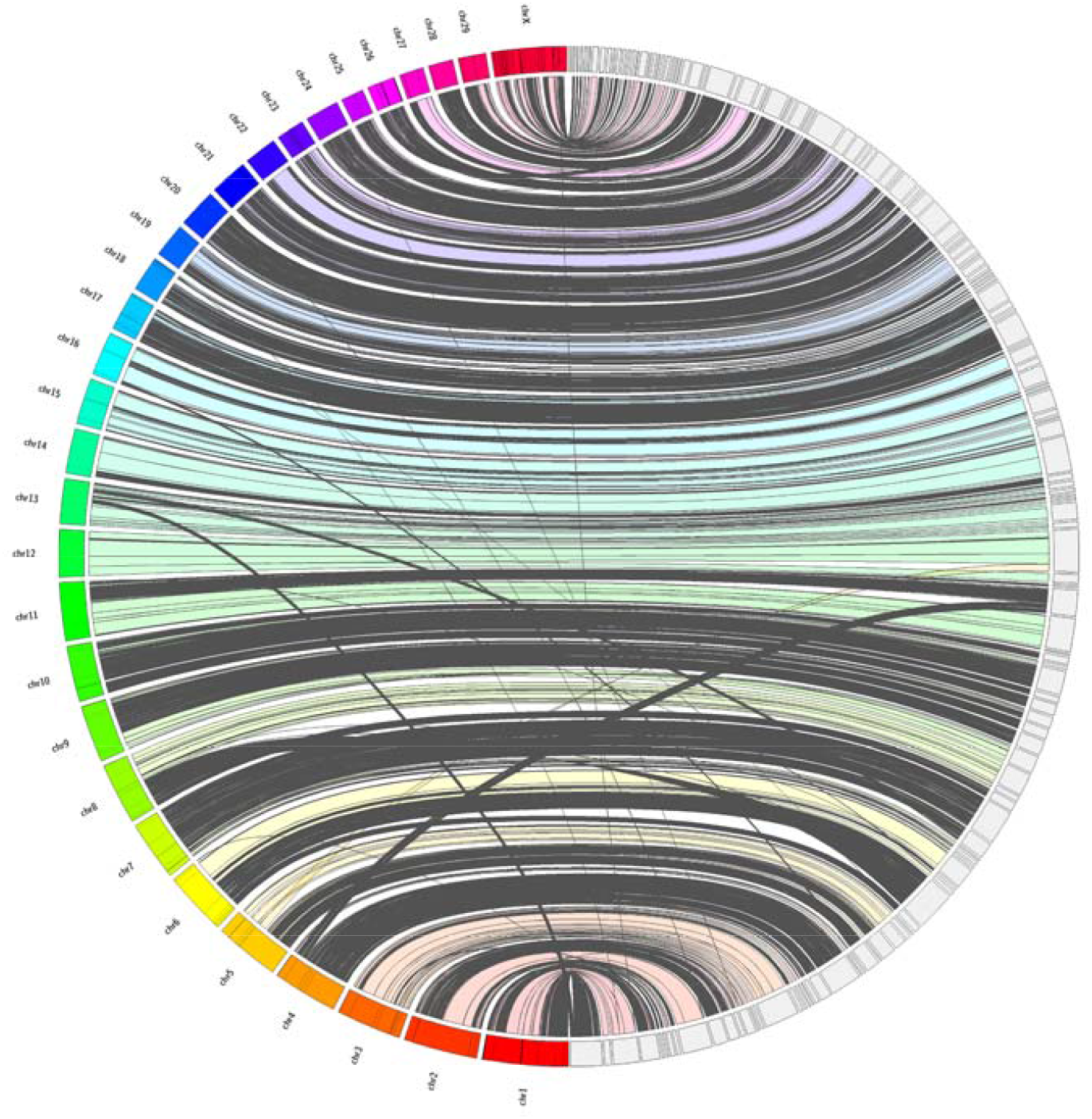
Synteny between caribou scaffolds and bovine chromosomes. *Rangifer tarandus* genomic scaffolds were aligned with the *Bos taurus* reference (ARS-UCD1.2) using JupiterPlot (https://github.com/JustinChu/JupiterPlot). Bovine chromosomes are labeled on the left and the 144 largest matching caribou scaffolds are represented on the right. Colored bands indicate syntenic regions in the same sense while grey bands indicate antisense synteny. Intersecting bands indicate non-syntenic regions between genome assemblies.

A phylogenetic tree rooted with the human genome was obtained using the single-copy orthologous genes from the *mammalia_odb10* database (Fig.4). In each of 10 species, 5,156 complete conserved genes were found and used to build the tree. As expected, caribou first clustered with mule deer (*Odocoileus hemionus*), a deer species common in western North America, and moose (*Alces alces*), another cervine inhabitant of the boreal forest. Together, these species represent the *Cervidae* clade and clustered with other Artiodactyla species including the *Bovidae* clade (including *Bos taurus, Bos indicus* and *Capra hircus*), *Suidae* (*Sus scrofa*) and *Camelidae* (*Camelus dromedarius*).

**Figure 4.**
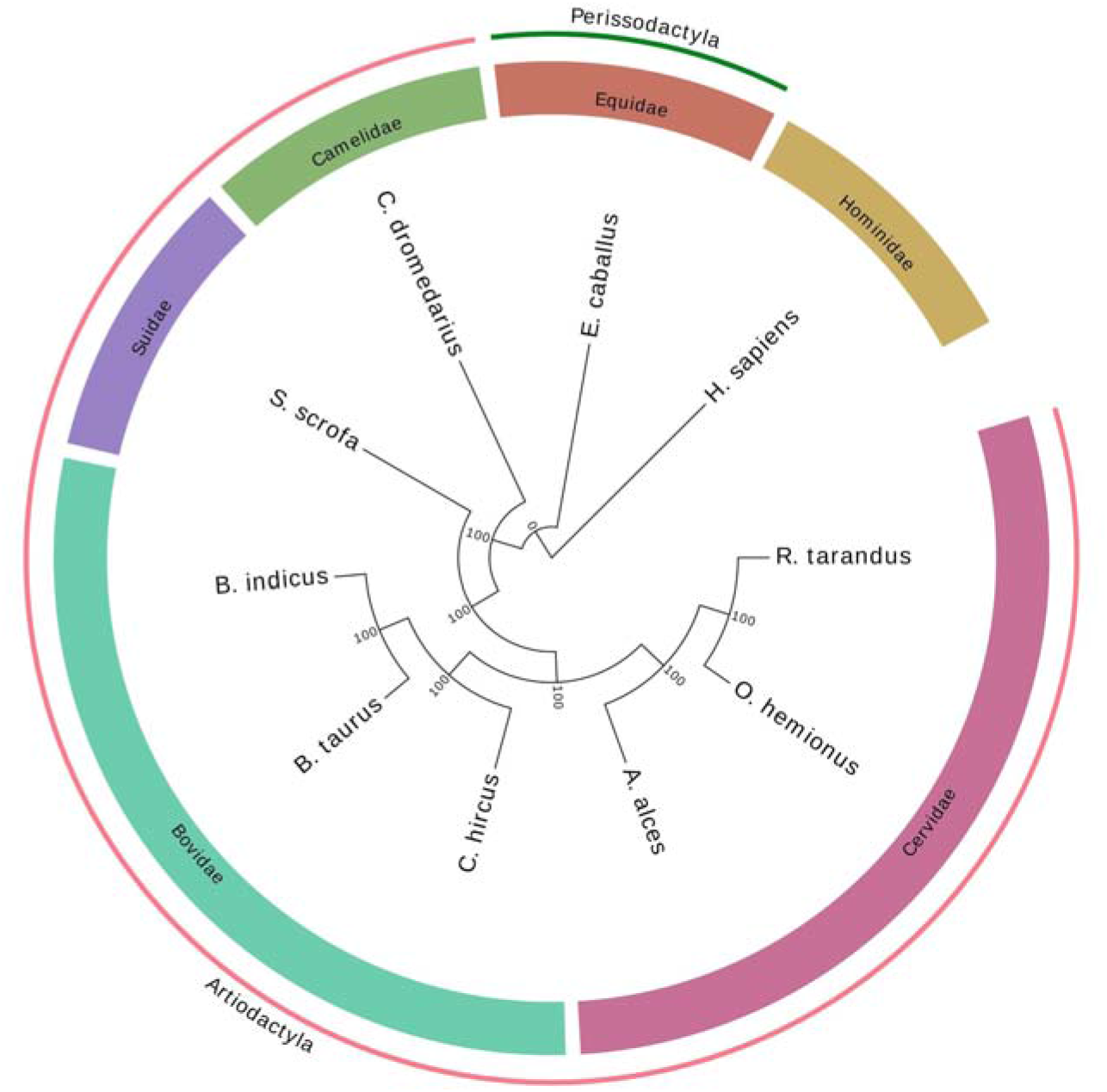
Phylogenetic tree of *Rangifer tarandus* and 9 other species based on the 5,156 complete orthologous genes, rooted using the human species.

### Genome annotation inferred from RNAseq de novo assembly

Gene expression diversity was maximized for annotation purposes by sequencing RNA extracted from several tissues (liver, muscle, blood, heart, lung, kidney, ovary). Read sequences were *de novo* assembled into transcripts using the *Transabyss* and *a5* bioinformatic tools, and then mapped on the genome assembly to identify coding regions (Fig. 1), which were annotated according to similarity with sequences in a Uniprot database. Since the complete annotated genomes closest to *Rangifer tarandus*, namely *Bos taurus* and *Capra hircus*, were annotated with putative functions based on similarity with the human genome, the cleaned Swissprot database (which includes only human sequences) was used (https://www.uniprot.org/proteomes/UP000005640) to avoid redundancy.

Transcripts were more numerous in the *Transabyss* transcriptome assembly (1,711,588) than in *a5* one (223,597). This process resulted in the identification of 17,172 annotated genes based on the *a5* assembly and 30,731 based on the *Transabyss* assembly. Overlap between both assemblies resulted in 20,419 corroborated annotated gene structures that were distributed over 2,759 genome assembly scaffolds. Among these, 3,025 coding sequences were annotated for transposable elements resulting in 17,394 gene models (gff3 file, Supplemental Data S1). Short coding sequences (< 500 bp) with low coverage (< 80%) or without homology with human gene sequences were not annotated.

### Large CNVs clustered in hotspots and encompassed coding sequences

Copy number variations were detected in 20 individuals representing the three *R. tarandus* ecotypes using 2^nd^-generation sequencing data and three types of evidence as implemented in the SpeedSeq tools suite (Chiang et al. 2015). Since our primary goal was to identify CNVs with adaptive potential, and thus subject to natural selection, rather than *de novo* CNVs not transmitted over generations, those detected in only one individual (or only in the reference assembly) were discarded. A total of 1,698 CNVs longer than 1,000 bp were detected over all samples, average length being 200,521 bp. The number of scaffolds containing at least one CNV was 162, and larger scaffolds contained more (supplemental Fig. S1). Altogether, CNVs accounted for 11.3% of the genome assembly (340,590,909 bp). Deletions were more numerous than duplications (1,466 versus 232) but significantly smaller (*t* = 3.7, p = 0.0002, supplemental Fig. S2). The number of CNVs per individual averaged 1,344.21 and ranged from 740 to 2,252 (Supplemental Fig. S3), while the average CNV locus frequency was 0.355. CNVs were not randomly distributed over the genome assembly but clustered into 31 hotspots including 227 CNVs (KS test; D = 0.047 and p = 0.001; Fig. 5). The number of CNVs per hotspot averaged 7.32 and reached 14. No scaffold contained more than 3 hotspots of CNVs.

**Figure 5:**
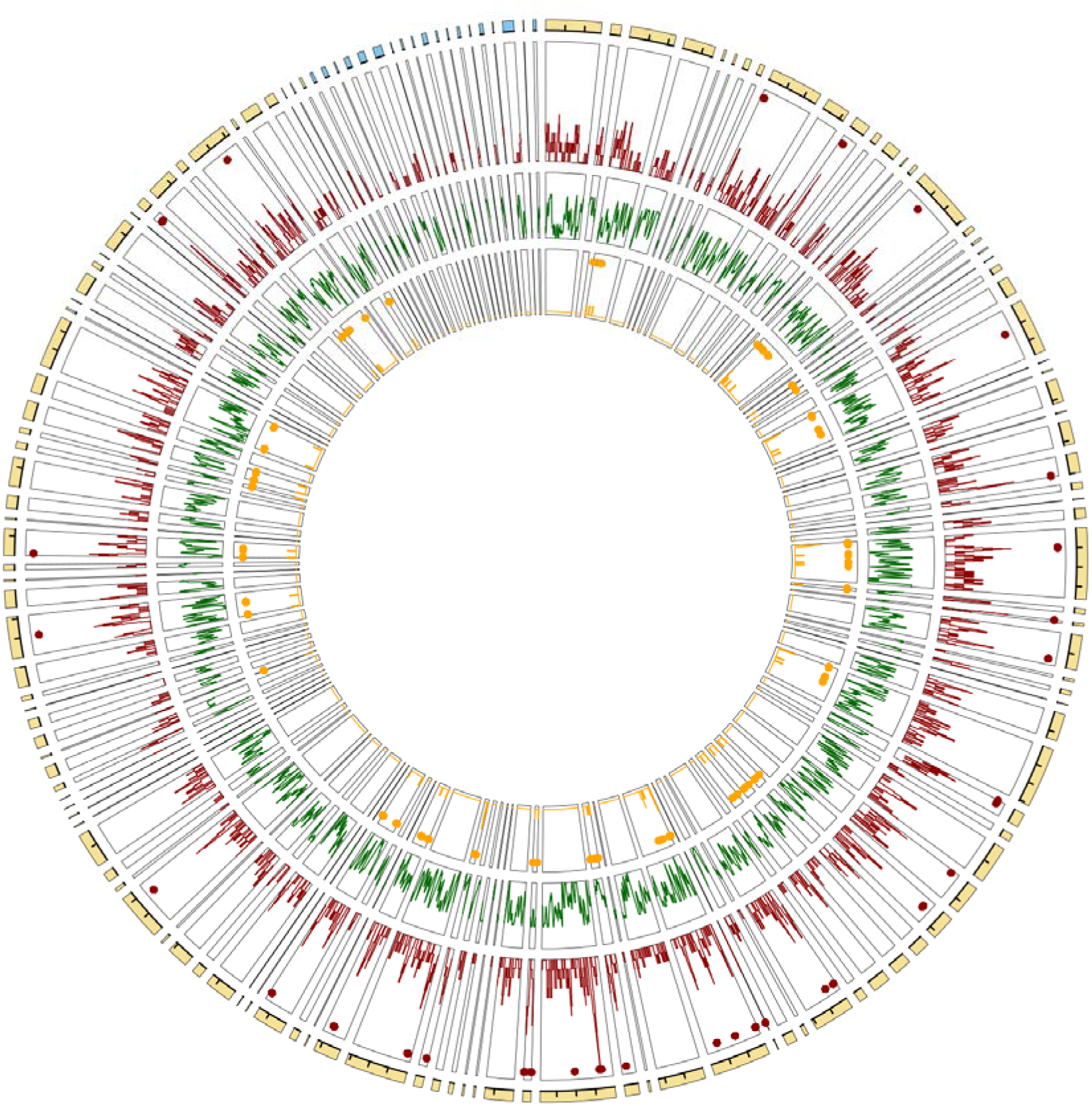
Genome architecture of CNVs, gene models and adaptive CNVs over the largest scaffolds in the new caribou genome assembly. From outward to inward track: the *Rangifer tarandus* new genome assembly with scaffolds matching autosomes (yellow) and the X chromosome (blue) in the *Bos taurus* assembly (interval between ticks = 20 Mbp), CNV density distribution (red) with hotspots marked as red dots, gene model density distribution (green), and distribution of likely adaptive CNVs (orange). The scaffolds are sorted according to the synteny with the *Bos taurus* genome.

A total of 332 of these large CNVs (19.5%) overlapped coding sequences, involving a total of 1,217 of the gene models identified in our genome assembly annotation. Duplications involved an average higher number of gene models (mean = 0.22 coding sequences per CNV, from 0 to 4) than deletions (mean = 0.20, from 0 to 5). The gene models involved in CNVs were annotated for functions altogether related to a large diversity of processes. An enrichment analysis in GO terms was performed and revealed a significant enrichment (adjusted p < 0.05) in various biological processes, including functions related to “regulation of protein metabolic process” (GO:0032269), “leukocyte activation” (GO:0045321), “muscle structure development” (GO:0061061), or “inflammatory response” (GO:0061061), amongst others (Supplemental Table S2).

Discriminant analysis of principal components (DAPC) was performed using the ‘adegenet’ R-package to identify CNVs for which the genetic distance between boreal sedentary and migrating ecotypes was maximal (Fig. 6); the mountain ecotype was not included because it was represented by a single individual. This revealed 43 CNVs showing 2.75% of the highest loading scores on the first principal component (Fig. 6B). Although most of these CNVs did not include any sequence annotated in our assembly, 15 were interestingly annotated for functions related to muscle and cardiac physiology, such as “musculoskeletal movement” and “regulation of heart rate”, temperature responses (“response to cold”), immune responses (“innate immune response” and “defense response to bacterium”), and environmental perception (“sensory perception of sound” and “visual perception” (Fig. 6C).

**Figure 6:**
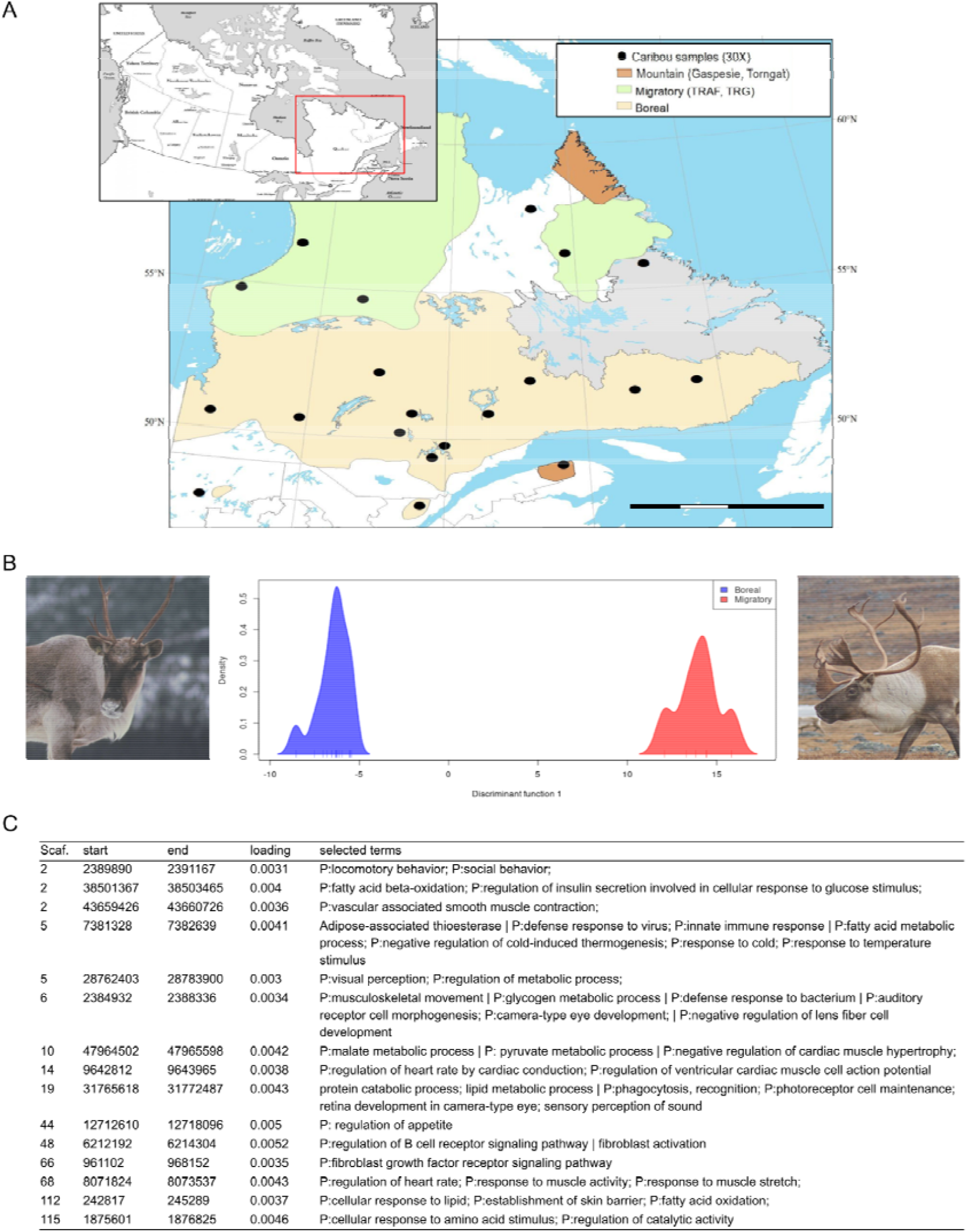
Divergent CNVs between caribou ecotypes in North-East America. (A) Ecotypes distribution and geographic locations for the 20 individuals sampled and sequenced (30X) for CNV detection; (B) Density distribution of the boreal sedentary (on the left) and migratory (on the right) caribou over the first axis of the DAPC based on CNVs; (C) Adaptation-related annotations for putative genes overlapping divergent CNVs.

## Discussion

### Genome assembly using different technologies

Each sequencing technology has its relative strength for *de novo* assembly of large genomes from non-model species. Whereas Illumina allows efficient sequencing of billions of high quality reads, these tend to remain short (< 1 kb), making it difficult to scaffold and improve large genome assembly contiguity (Coombe et al. 2018; R. L. Warren et al. 2015). However, linked reads corresponding to long DNA molecules of known origin (Chromium 10X), large insert sizes, or reads integrating remote DNA subsequences (DoveTail) allow scaffolding of short contigs into longer scaffolds that are informative of the DNA sequence distribution over the genome despite the occurrence of possibly large gaps. On the other hand, long-read sequencing can yield genome assemblies with higher contiguity, although reads are usually less numerous and of lower quality. To take advantage of each technology, our sequencing used short reads (Illumina), long reads (PacBio SMRT) and linked reads (Chromium 10X) assembled independently, each strategy yielding a genome assembly. These assemblies were then scaffolded using one another and public data to obtain the best genome assembly available to date for this species according to contiguity and correctness measures, for example, L90 = 131 (Table 1) while L90 = 289 in Taylor et al. (2019). However, short reads obtained from a 400 bp insert library did not allow us to improve our assembly, which was unexpected in view of previous findings (Jackman et al. 2018). This was likely due to the very high yield (107X) that we obtained for linked reads that were also short reads with the same very low sequencing error rate. Linked-read sequencing thus proved to be a very interesting strategy for *de novo* assembly of a large genome.

In terms of contiguity and accuracy (BUSCO analysis; Table 1), this new genome assembly compares very well with other recent genome assemblies for livestock species such as chicken (W. C. Warren et al. 2017) or other wild species such as the grizzly bear (Taylor et al. 2018) or sea otter (Jones et al. 2017) and was superior to those of other *Cervidae* species (Dussex et al. 2020; Upadhyay et al. 2020). Notably, the highest quality genome assemblies (including the present one) are usually obtained when different sequencing strategies are used, including long reads and linked reads, often in combination with typical short-read sequencing (Wallberg et al. 2019; Kongsstovu et al. 2019).

Consistent with the high number of orthologous genes found in our new caribou genome assembly, the phylogenetic tree obtained using those sequences (Fig. 4) presented the expected species relationships, with *Bovidae* being the clade closest to *Cervidae*, which included moose and mule deer. These two families have several characteristics in common (two-toed ungulates, ruminants), including a similar genome size and overall structure inherited from a common ancestor. However, fissions of 6 chromosomes changed the number of chromosomes from 29 to 35 in *Cervidae*, while a fission of chromosomes 26 and 28 brought the total to 30 in *Bovidae* (Frohlich et al. 2017). Scaffolds from *Cervidae* genome assemblies therefore show a high synteny with the cow reference genome (Taylor et al. 2019; Li et al. 2017; Bana et al. 2018) although many scaffolds should map to the chromosomes that were split (1, 2, 5, 6, 8 and 9) in the course of genome evolution since the last common ancestor. A caribou scaffold might likewise overlap bovine chromosomes 26 and 28, since these two should form only one chromosome in *Cervidae*. The genome comparison illustrated by the Jupiter plot (Fig. 3) indicated very high synteny with only 18 bands and lines illustrating variations in the DNA segment order. One of these crossing bands is a caribou scaffold that maps partially to bovine chromosomes 26 and 28. The remaining crossing lines and bands are indicative of either chimeric assemblies or translocations. *In situ* DNA sequence marking and microscopic visualization such as fluorescence in situ hybridization (FISH) would undoubtedly help to resolve such uncertainty. Nevertheless, so few discrepancies (excluding the expected one) between bovine and caribou genome assemblies compared to previous reports illustrates the genome assembly improvements. In addition, clustering the largest scaffolds into chromosomes using FISH, for instance, is now possible, given the relatively low number of scaffolds representing 90% of the genome assembly.

This contiguous and accurate assembly will undoubtedly pave the way to other genomic tool developments and genomic investigations of this threatened species, such as landscape genomics and genomics of adaptation at the population level.

### Genome annotation based on expressed sequences

To this end, another key aspect of genomic investigations is genome assembly annotation. Despite much progress in recent years, annotation of a genome based on DNA motifs and gene prediction remains challenging and time-consuming and needs constant updating (Salzberg 2019). As a first step towards this goal, we used RNAseq of a composite sample representing several tissues to assemble a transcriptome with high diversity allowing identification of thousands of transcript sequences distributed throughout the genome. Identity with known proteins in the Uniprot database allowed annotation of a large subset of these transcripts with putative functions.

The RNAseq based approach is very interesting since it allows identifying genome regions that are truly transcribed, thus avoiding most of the issues related to the occurrence of unexpressed pseudogenes and spurious identification of non-genes when predicting directly from the genome assembly. We took full advantage of this feature by discarding transcripts that could not be annotated due to lack of homology with known coding sequences in the human genome. These transcripts were often short and possibly represented pseudogenes with incomplete reading frames. However, such an RNAseq approach requires analyzing the greatest possible diversity of samples in terms of tissue, environmental conditions, time points (circadian variation) and developmental stage (embryo, juvenile and adult of both sexes) to obtain an exhaustive annotation of the genome. Given the ever-increasing affordability of sequencing, this could be achievable in the foreseeable future. Nevertheless, much of the gene models set was likely reported here, given that 17,394 gene models were identified, which would represent 79% of the complete set assuming a number of genes similar to the one of *Bos taurus*, for which the most recent annotation includes 21,880 gene models (ensembl, release104).

Because of the relative proximity to the most intensely studied mammalian models (cow, mouse, rat, human), our annotation of coding sequences based on identity with known sequences was successful overall, with few unknown functions. However, Gene Ontology terms enrichment analyses are based on reported gene functions, which are currently associated mostly with human pathologies and disorders. Far fewer annotations relate to responses to natural environmental pressures into the wild. It is therefore possible that annotations relevant to differentiation between ecotypes (adaptations) were hidden in an excessive amount of annotations related to human pathologies (cancer or neurocerebral issues for example). Efforts to characterize the coding sequence molecular functions and gene ontology annotations with regards to natural environmental conditions would be beneficial to future studies focused on genetic variations in a wildlife conservation context.

### Hotspots of CNVs detected in wild mammals

Genomes of domesticated mammals (including ruminants) have been entirely sequenced and studied intensively for decades, leading to the development of SNP or aCGH chips used to characterize many individuals. As methods and softwares making use of these resources to detect CNVs were developed, those chips have been largely used to detect such variations in a number of species, races, and lineages (Clop, Vidal, and Amills 2012). However, few early chips were of sufficient density to cover entire genomes (Carvalho et al. 2004) and additional CNVs were discovered when whole genome sequencing (WGS) became widespread (Alkan, Coe, and Eichler 2011). Our WGS data revealed CNVs in 20 individuals from different ecotypes and geographic origins. As expected, we found a number of large CNVs (size > 1,000 bp) in the same range of numbers reported in previous CNV studies using the same detection approaches (Bickhart et al. 2012; Paudel et al. 2015; Schurink et al. 2018) and far more than historically detected in domesticated mammals using SNPchips and aCGH (Clop, Vidal, and Amills 2012). These long CNVs covered 11.3% of the genome assembly, as observed for other species such as human (11-12%, Stankiewicz and Lupski 2010; Redon et al. 2006) and horses (11.2%, Schurink et al. 2018; Ghosh et al. 2014) using similar detection parameters.

These large CNVs were distributed throughout the caribou genome assembly with hotspots including up to 14 CNVs. This genome architecture of CNVs including hotspots is widespread among living organisms and has been observed not only in humans and chimpanzees (Perry et al. 2006) and other mammals (Yang et al. 2018; Clop, Vidal and Amills 2012) but also a wide range of plants (Muñoz-Amatriaín et al. 2013; Torkamaneh et al. 2018; Prunier et al. 2019; Swanson-Wagner et al. 2010). This universality is mainly explained by the main molecular mechanisms that lead to the formation of large CNVs, which are related to the occurrence of tandem repeats (Hastings et al. 2009; Perry et al. 2006). This CNVs genome distribution including hotspots is not trivial in evolutionary terms since advantageous copy numbers are likely to aggregate into heritable clusters (Yeaman 2013). This trend may even be amplified since CNVs may prevent recombination and thus favor the persistence of large genomic islands of divergence (Tigano et al., n.d.).

Another feature usually observed in whole genome scans for CNVs is the higher number of deletions than duplications. This has long been attributed to a detection bias associated with SNP chips or aCGH, which are more prone to identify deletions that result in 2-fold variations than duplications that result in 1.5-fold variations in diploid genomes (Carter 2007; Alkan, Coe, and Eichler 2011). However, this should not affect CNV detection based on sequencing data as much, since coverage is only one element taken into account to identify a CNV (Chiang et al. 2015) and there is no obvious reason why split-read-based or split-read-pair-based detection would be biased towards deletions. Consistent with this, a number of recent sequencing-data-based studies show similar numbers of duplications and deletions (Sudmant et al. 2015; Zheng et al. 2016) although the number of detected CNVs is much higher and down to 200 bp versus 1 kb in these earlier studies, thus limiting comparability. Another possible factor contributing to higher numbers of deletions than duplications among large CNVs is one of the mechanisms leading to CNVs that results in the loss of DNA segments, namely the intra-chromatid NAHR (Gu, Zhang, and Lupski 2008). The prevalence of this mechanism has not been demonstrated to our knowledge but the higher proportion of large deletions detected in the caribou genome (86%) suggests that it may be considerable. This finds support in another sequencing-data-based CNV study of cats in which the prevalence of losses was 84% using a detection threshold of 5 kb for CNV length (Genova et al. 2018). In addition, the prevalence of deletions was 90% in a recent report on dogs using a CNV minimal size of 1 kb (Serres-Armero et al. 2021). Intra-chromatid NAHR thus appears to contribute to long DNA segment deletions, of which the signal is blurred by other mechanisms when shorter CNVs are included. Meta-analysis of proportions of deletions and duplications in different CNV length ranges in a variety of species would settle this question and more generally help classify structural variations that occur over a broad range of DNA lengths, from small indels to chromosomal rearrangements (Mérot et al. 2020).

Despite this new assembly representing 86–89% of the entire genome and the number of CNVs being close to those reported for other mammals (Yang et al. 2018) suggesting that we gathered a major proportion of the common CNVs, additional CNVs may occur in other ecotypes or in other parts of the species distribution. The number of detected CNVs is a measure of the CNV genetic diversity and is subject to the same detection parameters and evolutionary forces as the genetic diversity of any polymorphism. First, genetic polymorphisms are usually found in higher numbers when more individuals are studied, and CNV diversity is strongly related to the number of tested individuals (Bickhart et al. 2012; Conrad et al. 2010). Second, a hierarchical population structure is expected at the entire species distribution level with some CNVs being peculiar to specific populations (Conrad et al. 2010; Sudmant et al. 2015) or lineages (Hu et al. 2020; Yang et al. 2018). Alleles thus remain undetected when testing individuals from a fraction of the species range. Third, like SNPs, rare CNVs can be peculiar to one individual. Testing 20 individuals from a subpart of the species distribution possibly limited our detection power. However, since CNVs present higher mutation rates than SNPs (Lupski 2007), rare alleles in CNVs possibly result from *de novo* formation limited to the sampled tissue and have not likely spread into the germline. Such rare CNVs provide little insight into adaptive evolution in wild species and were not targeted in this study. By sampling various ecotypes and geographic origins, we likely increased the CNV diversity and the odds of detecting CNVs related to adaptation beyond the limits of the sampled area.

### CNVs signatures related to adaptation in wild mammal ecotypes

Lengthy CNVs may span entire gene-coding sequences and lead to gene expression variations, or partially overlap gene sequence, thus disrupting transcript sequence with variable phenotypic impacts (Lupski and Stankiewicz 2005). In any case, CNVs that include coding sequences are more likely than intergenic CNVs to have such impact, since the involvement of gene copy number in phenotypic variation is reported widely. One example in humans is starch-digesting ability, proportional to the number of copies of the *AMY1* gene, which encodes salivary amylase (Perry et al. 2007). Similarly, farm animal coat color is often associated with gene copy number variations (Clop, Vidal, and Amills 2012), for example, the *ASIP* gene for light pigmentation in sheep (Dong et al. 2015). Based on annotation of our caribou genome assembly, 19.5% of the CNVs overlap with gene model sequences. Annotations of these gene models represented a large diversity of biological processes enriched in GO terms related to immunity and healing, metabolism, musculoskeletal development, or environmental perception, amongst others (supplemental Table S2). Most of these terms have been revealed in previous enrichment analyses of genes in CNVs in mammals, such as metabolism and olfactory perception in swine (Paudel et al. 2015), immune responses in chimpanzees (Perry et al. 2006), horses (Schurink et al. 2018) and other farm animals (Clop, Vidal, and Amills 2012), cardiac and skeletal muscles in humans (Conrad et al. 2010), and body height in dogs (Serres-Armero et al. 2021). The phenotypic variations associated with CNV diversity in model organisms present a high adaptive potential for wild species such as caribou. However, the genome annotation was based on expressed sequences in a multi-tissue pooled sample. Genes not expressed in these tissues under these conditions were missed, making the list of gene models incomplete. The CNVs may encompass additional coding sequences not described here, although current annotations of identified gene models included in CNVs support their potential involvement in adaptation.

In our comparison of sedentary and migrating caribou, the DAPC analysis revealed 43 CNVs that contributed the most to the variability and are thus promising candidates to adaptive divergence. Fifteen of these overlapped gene sequences that were annotated for relevant biological processes (Fig. 6). First, it is well known that migrating caribou roam hundreds to thousands of kilometers annually while sedentary (boreal) ones travel much less (Mallory and Hillis 1998). Annotations related to “muscle contraction”, “heart development”, “cardiac muscle hypertrophy” and “cardiac muscle contraction”, as well as “musculoskeletal movement” and “locomotory behavior” were therefore unsurprising and consistent with this difference in habitat range. Similarly, annotations related to “fat metabolism”, “response to cold”, or “vascular associated smooth muscle contraction” are consistent with the summer temperature differences between the tundra and the boreal forest and with the particular fat metabolism and heat loss mitigation by peripheral vasoconstriction reported in this species (Blix 2016). Furthermore, five CNVs included genes annotated for functions related to “defense responses” and “immunity”, including “skin barrier” annotation. As migrating caribou reaching the tundra are harassed during the summer by Oestridae parasitic flies that lay eggs under their skin (Hagemoen and Reimers 2002) while sedentary caribou in the boreal forest are relatively spared by these flies, some CNV diversity between ecotypes was to be expected. Other interesting annotations included “sensory perception of sound”, “visual perception” and “retina development”. Given that summer habitats of migrating and sedentary caribou differ considerably in terms of forest canopy, these terms are likely related to adaptation to local conditions where sight or sense of hearing may be differentially favored. We also noted the terms “social behavior” and “regulation of appetite” which may be related to the differential group composition and access to summer forage. While sedentary caribou form small groups and have access to small patches of edible vegetation spread regularly throughout the boreal forest, migrating caribou travel in large herds for kilometers to reach large patches of edible vegetation where intra-specific competition can be important, thus alternating between dietary abundance and scarcity.

We also noted terms with slightly lower loading scores but interesting from the perspective of adaptation and knowledge acquired from the study of Eurasian reindeer. These terms referred to light and circadian cycles such as “response to UV” (N = 44), “regulation of circadian sleep/wake cycle” (N = 2) and “vitamin-D”-related annotations (N = 15). In the spring, migrating caribou travel north, closer to the Arctic Circle, where summer nights are shorter than in the boreal forest. Thus, migrating caribous likely manage active and resting periods differently than sedentary caribou. It has been shown in the european reindeer that melatonin secretion in reindeer is highly sensitive to ambient light rather than regulated by an internal circadian clock (Stokkan et al. 2007) and more importantly, that such differences in day/night activity cycles exist between two *Rangifer tarandus* subspecies, one inhabiting latitudes north of the arctic circle (Svalbard, Norway) and the other inhabiting northern Europe (mainland Norway) (van Oort et al. 2005). In addition to this differential exposure to daylight, canopy opening also contributes to UV exposure, making “response to UV” an expected annotation.

Altogether these promising annotations for genes included in CNVs points toward a role of CNVs in adaptation to local conditions in wild species. While CNVs including gene models with such annotations are more interesting, the possibility of CNVs affecting gene expression, by influencing promoters or through positional effects (Lupski and Stankiewicz 2005), should not be overlooked, since these can have relevant physiological implications. Thus, further investigation of the functional aspects of all CNVs may be of interest though represent a daunting task. Nevertheless, these CNVs including genes annotated for functions potentially linked to ecotype divergent adaptive traits appeared worth being tested in large populations. Indeed, comparing 19 individuals, as presented here, is a very important first step towards the identification of adaptive CNVs but testing a non-random distribution over populations and ecotypes would further support the involvement of the CNVs in phenotypic variations in response to selective pressure (see Serres-Armero et al. 2021 for an example in dogs). Since CNV detection requires relatively extensive sequencing (Lupski and Stankiewicz 2005; Layer et al. 2014), testing several individuals with focus on the candidate CNVs reported here should allow evaluation of their impact on phenotypes and adaptations.

## Conclusions

*De novo* assembly of large genomes is a difficult undertaking, particularly for undomesticated species, which usually present less economical interest and are consequently not, or less, described at the genome level. Genome contiguity may be reached at the expense of accuracy, although both objectives are attainable using recently developed long-read sequencing technologies. In this study, we built a new genome assembly (JAHWTM000000000, Bioproject: PRJA739179) made mainly of a few large scaffolds that allowed the first genome architecture analysis in this species including gene models and CNVs. RNA sequencing allowed us to publicly release a first robust genome annotation for *Rangifer tarandus* (Supplemental Data S1), which will undoubtedly pave the way to the development of genomics tools such as SNP-based genotyping chips, allowing to inform species conservation and management efforts for this species. Detecting CNVs between migrating and sedentary caribou ecotypes yielded a list of CNVs encompassing annotated genes that imply a role for CNVs in adaptation of this northern wild ruminant.

## Materials and Methods

### Whole genome long-read sequencing

Previous caribou/reindeer assemblies were made using blood as the source of DNA (Li et al. 2017; Weldenegodguad et al. 2020), which is known to hinder genome assembly (Rosen et al. 2020). In addition, two interesting sequencing technologies that could improve genome assembly contiguity have not been used to date to assemble the *Rangifer tarandus* genome, namely Pacific Biosciences and 10XGenomics technologies.

Since the single-molecule real-time (SMRT) technique (Pacific Biosciences, Menlo Park, CA, USA) does not require DNA amplification prior to sequencing and results depend largely on DNA initial quality, high-molecular-mass (100–200 kbp) genomic DNA from muscle biopsy was isolated using a MagAttract HMW kit according to the manufacturer’s instructions (QIAgen). DNA quantity and quality were evaluated on genomic DNA screentape using a 4200 Tapestation (Agilent) and retaining only peaks of mass > 45 kbp. The library was prepared and SMRT sequencing (24 runs aiming for 30X coverage, 4 Gb of data per SMRT cell) was performed on the Sequel machine at Génome Québec (Center of Expertise and Services, Montréal, QC, Canada).

To reconstruct long DNA fragments, linked-read sequencing was also performed. Chromium 10X libraries (from 10XGenomics) were prepared at Génome Québec using the same high molecular weight genomic DNA as for SMRT sequencing. Paired end (150 bp) sequencing was performed on an Illumina HiSeqX (at Génome Québec Center of Expertise and Services, Montréal, QC, Canada). Three sequencing lanes were run to obtain approximately 100X genome coverage.

### Transcriptome analyses

A pool of mixed samples (including liver, muscle, blood, heart, lung, kidney and ovary) was collected and transported in RNAlater stabilization solution (ThermoFisher, Waltham, MA, USA) and stored at −20°C until RNA extraction. RNA isolation was performed using TRIzol reagent (ThermoFisher) as per the manufacturer’s RNA isolation protocol, followed by on-column purification and DNAse I treatment (PicoPure, ThermoFisher). RNA quality and integrity were assessed using RNA screentape on a 4200 TapeStation system (Agilent, Santa Clara, CA, USA). Only RNA with an integrity number over 7 was used for library preparation and sequencing.

Transcriptomes were sequenced using paired-end 150 bp Illumina HiSeq X (Illumina, San Diego, CA, USA) at Génome Québec Center of Expertise and Services (Montréal, QC, Canada) with NEB mRNA stranded Library preparation (New England Biolabs, Whitby, ON, Canada).

### Whole-genome short-read sequencing of the various ecotypes

Ear punch flesh was collected from 20 individuals (10 females and 10 males) in different regions of the Province of Québec (Canada) to include migratory, sedentary (boreal) and mountain ecotypes. Genomic DNA was isolated from frozen ear punches using DNeasy Blood and Tissue kits (QIAgen, Toronto, ON, Canada). DNA quantity and integrity were evaluated using genomic DNA screentape on a 4200 TapeStation system (Agilent). Only samples with a DNA integrity superior to 7 were used. Shotgun sequencing was performed using a PCR-free DNA library preparation (NEBNext Ultra II DNA Library Prep Kit, New England Biolabs). Libraries were paired end 150 bp sequenced with Illumina HiSeqX. A genome coverage of approximately 30X was obtained from 20 lanes of sequencing.

### Bioinformatics analyses

#### Genome Assembly

The genome assembly was built from three approaches based on the three different sequence data types (Fig. 1). First, high-quality long reads from PacBio sequencing were selected and assembled using the Falcon assembler v.1.4.2 (Chin et al. 2016, 2013). This assembler aligns autocorrected long reads to each other and assembles these into contigs. Then linked reads obtained from Chromium 10X sequencing were assembled independently using the Supernova assembler (Zheng et al. 2016; Marks et al. 2019; Weisenfeld et al. 2017). This assembler is an adapted version of DISCOVAR, an assembler designed to assemble short reads using De Debruijn graphs (Weisenfeld et al. 2014), that takes into account barcodes to pair reads and thereby elongate contigs and scaffolds. Finally, the short reads from the individual with the highest coverage among the 20 individuals were assembled using DISCOVAR-denovo, an assembler optimized to assemble genomes with size close to 3 Gb from high-quality short reads.

The Falcon assembly was scaffolded using the Supernova assembly and LINKS (Warren et al. 2015) to yield a second assembly. This second assembly was scaffolded again using the same bioinformatic tool and the publicly available genome assembly based on DoveTail sequencing (Taylor et al. 2019).

#### Annotation based on transcriptome assembly from RNAseq data

##### RNA assemblies

Read quality was assessed using Fastqc and reads were then cleaned using Trimmomatic v0.36 (Bolger, Lohse, and Usadel 2014). Cleaned reads were assembled twice using the SGA (Simpson and Durbin 2012) and IDBA-UD assemblers (Peng et al. 2012) via the a5 perl pipeline (Coil, Jospin, and Darling 2015) and the Trans-ABySS assembler (Robertson et al. 2010). Both assemblies were kept for the next step since these algorithms may assemble RNA differently (e.g., more contiguously or less so) while pointing to the same gene regions.

##### GAWN

The two transcriptome assemblies were then used to annotate the genome assembly using the GAWN pipeline (https://github.coverm/enormandeau/gawn) that maps transcriptome sequences onto the genome assembly using GMAP (Wu and Watanabe 2005) to produce a gff3 file and gathers annotations from the Swissprot database (UniProt Consortium 2019) using BLASTX (Altschul et al. 1990). Overlapping gene structures found in both transcriptomes using in-house scripts and the “merge” function from the bedtools suite (Quinlan and Hall 2010) were deemed more reliable and thus included in the final annotation file (Supplemental Data S1).

##### Phylogeny

Single-copy orthologous genes from mammalia_odb10 found using BUSCO v3.0.2 (Waterhouse et al. 2018; Simão et al. 2015) with lineage dataset for 10 species including *Homo sapiens* as an outgroup were used for phylogenetic analysis. Common single-copy-gene DNA sequences were aligned using MAFFT v7.397 (Katoh and Standley 2013) and trimmed using trimAl v1.4 (Capella-Gutiérrez, Silla-Martínez, and Gabaldón 2009). Gene sequences were then concatenated to form a single sequence per species. The phylogenetic tree was inferred using RAxML v8.2.11 (Stamatakis 2014) with the GTR+I+G substitution model previously selected by JModelTest v2.1.10 (Darriba et al. 2012).

##### CNV detection

Structural variations were detected using the SpeedSeq tools suite (Chiang et al. 2015). Paired-end reads obtained from the 20 individuals were first cleaned using Trimmomatic v0.36 (Bolger, Lohse, and Usadel 2014) and aligned to our newly built genome assembly using “speedseq align”. SNVs were then detected independently for each individual using “speedseq sv”, which runs LUMPY (Layer et al. 2014). LUMPY uses three types of evidence to declare an SNV, namely read pairs, split reads and generic read depth (in our case using CNVnator (Abyzov et al. 2011) optional analysis). All detected structural variations were then concatenated, and all samples were genotyped for these variations using “svtyper” (https://github.com/hall-lab/svtyper). Variations occurring within only one genome were excluded, since they were deemed less reliable and may have been the result of *de novo* tissue-specific CNVs not transmitted over generations.

The non-random CNV distribution was tested using a genome-wide KS-test between the distributions of non-CNV and CNV positions. In addition, sliding window analysis was performed to identify CNV hotspots based on the average number of CNVs within 2 Mb windows (pace 1kb) and regions constituted of contiguous windows with average in the higher tail (above 97.5%) of the distribution were deemed hotspots of CNVs.

## Acknowledgements

We thank the Centre d’Expertise et des Services Génome Québec at the Centre Hospitalier de Sainte-Justine, Montréal (QC, Canada) for their contribution to molecular data production. We also thank ComputeCanada and CalculQuébec for access to large computing resources. This project was funded through a project grant awarded to CR, SDC and AD in the Genomic Applications Partnership Program of Genome Canada. We acknowledge co-financing by Genome Quebec and the Ministère des Forêts, de la Faune et des Parcs du Québec.

## Declaration

All samples were collected from live captures as part of caribou population monitoring by the province of Quebec (Canada). Live capture and sample collection followed the Canadian Council on Animal Care guidelines, and all procedures were approved by Animal Care Committees (CPA-FAUNE 18-04). RNA samples were collected from an individual that died during handling.

## Supplemental material

Supplemental Data S1: A gff3 file including all coding sequences inferred from RNAseq de novo assembly

Supplemental Table S2: A table with Gene Ontology terms significantly enriched in all CNVs.

Supplemental Data S3: including supplemental figures Supp. Fig. S1, Supp. Fig. S2, and Supp. Fig. S3.

